# DciA helicase operators exhibit diversity across bacterial phyla

**DOI:** 10.1101/2022.01.24.477630

**Authors:** Helen C. Blaine, Joseph T. Burke, Janani Ravi, Christina L. Stallings

## Abstract

A fundamental requirement for life is the replication of an organism’s DNA. Studies in *Escherichia coli* and *Bacillus subtilis* have set the paradigm for DNA replication in bacteria. During replication initiation in *E. coli* and *B. subtilis*, the replicative helicase is loaded onto the DNA at the origin of replication by an ATPase helicase loader. However, most bacteria do not encode homologs to the helicase loaders in *E. coli* and *B. subtilis*. Recent work has identified the DciA protein as a predicted helicase operator that may perform a function analogous to the helicase loaders in *E. coli* and *B. subtilis*. DciA proteins, which are defined by the presence of a DUF721 domain (termed the DciA domain herein), are conserved in most bacteria but have only been studied in mycobacteria and γ-proteobacteria (*Pseudomonas aeruginosa* and *Vibrio cholerae*). Sequences outside of the DciA domain in *Mycobacterium tuberculosis* DciA are essential for protein function but are not conserved in the *P. aeruginosa* and *V. cholerae* homologs, raising questions regarding the conservation and evolution of DciA proteins across bacterial phyla. To comprehensively define the DciA protein family, we took a computational evolutionary approach and analyzed domain architectures and sequence properties of DciA-domain containing proteins across the tree of life. These analyses identified lineage-specific domain architectures amongst DciA homologs as well as broadly conserved sequence-structural motifs. The diversity of DciA proteins represents the evolution of helicase operation in bacterial DNA replication and highlights the need for phylum-specific analyses of this fundamental biological process.

**IMPORTANCE:** Despite the fundamental importance of DNA replication for life, this process remains understudied in bacteria outside of *Escherichia coli* and *Bacillus subtilis*. In particular, most bacteria do not encode the helicase loading proteins that are essential in *E. coli* and *B. subtilis* for DNA replication. Instead, most bacteria encode a DciA homolog that likely constitutes the predominant mechanism of helicase operation in bacteria. However, it is still unknown how DciA structure and function compare across diverse phyla that encode DciA proteins. In this study, we perform computational evolutionary analyses to uncover tremendous diversity amongst DciA homologs. These studies provide a significant advance in our understanding of an essential component of the bacterial DNA replication machinery.

## INTRODUCTION

DNA replication is a process critical to life for all organisms. The current paradigm for the process of DNA replication in bacteria has primarily been based on studies in *Escherichia coli* and *Bacillus subtilis*. Bacterial DNA replication begins with the binding of the replication initiation protein DnaA to specific sequences referred to as DnaA boxes at the origin of replication (oriC) (1-7). DnaA binding to double-stranded DNA (dsDNA) triggers DNA unwinding at an AT-rich region of DNA called the DNA unwinding element (DUE), leaving a bubble of single-stranded DNA (ssDNA) (1, 4, 6, 8, 9). The ssDNA bubble is coated by single-stranded binding protein (SSB) (10), followed by the concerted loading of two hexameric replicative helicases onto the SSB-coated replication fork. The two helicases translocate along the two sides of the replication fork, unwinding the dsDNA as they move (1, 4, 5, 11–14).

Bacterial replicative helicases (DnaB in *E. coli* and DnaC in *B. subtilis*) are Superfamily IV type helicases, which are defined as hexameric RecA ATPases (4,15,16) that translocate in the 5’-3’ direction (11,12,17) The bacterial replicative helicase translocates on ssDNA using a “hand-over-hand” mechanism, which is driven by nucleotide hydrolysis (18, 19) (reviewed in (17)). The C-terminus of the bacterial replicative helicase contains the RecA-like fold that is responsible for the ATPase activity and is connected to an N-terminal scaffolding domain via a linker region (1,20,21). The replicative helicase must oligomerize into a double-layered hexameric ring to be active during replication, with one layer made up of the N-termini and the other layer composed of the C-termini (1,22,23). In *E. coli* and *B. subtilis*, the loading of the replicative helicase is performed with the help of a helicase loader, termed DnaC in *E. coli* and DnaI in *B. subtilis* (4, 24–27). *dnaC* and *dnaI* were acquired by *E. coli* and *B. subtilis*, respectively, via domestication of related but distinct phage ATPase-containing genes (28-31). DnaC and DnaI are both in the ATPases Associated with diverse cellular Activities (AAA+) ATPase family, and the ATPase activity of DnaC is required for its helicase loading function at the origin of replication (32, 33).

*E. coli* and *B. subtilis* have long represented the paradigm of helicase loading during bacterial replication. However, most bacteria do not encode ATPase helicase loader homologs to DnaC or DnaI. Instead, most bacteria encode the ancestral protein, DciA (DnaC/I Antecedent) (28, 34), which is defined by the presence of a Domain of Unknown Function DUF721 (termed DciA domain herein). Despite the prevalence of DciA domain containing proteins in bacteria (28), DciA homologs have only been studied in actinobacterial (*Mycobacterium tuberculosis and Mycobacterium smegmatis)* and *γ-*proteobacterial (*Pseudomonas aeruginosa* and *Vibrio cholerae*) species (28, 34–36). DciA homologs interact with the replicative helicase DnaB and are essential for *M. tuberculosis, M. smegmatis,* and *P. aeruginosa* DNA replication and viability (28, 34–36). Based on DciA’s interaction with the replicative helicase and requirement for DNA replication, DciA has been proposed to perform a function analogous to that of the DnaC/I helicase loaders. However, DciA does not have a predicted ATPase domain and, therefore, cannot be considered a helicase loader like DnaC/I. Instead, DciA is referred to as a predicted helicase operator, although the mechanism of DciA helicase operation is still unknown (28, 34).

Based on studies in *M. tuberculosis*, the DciA domain was predicted to contain a region of structural homology to the N-terminus of DnaA (34), which was subsequently confirmed in *V. cholerae* DciA (35,36). The DciA domain in the *M. tuberculosis* DciA homolog is essential for protein function (34). In addition, mycobacterial DciA homologs encode a 58 amino acid (aa) sequence extension N-terminal to the DciA domain that is essential for *M. smegmatis* viability (34). However, this sequence is not conserved in *P. aeruginosa* or *V. cholerae* DciA. Instead, sequences C-terminal to the DciA domain in *V. cholerae* DciA that are not shared with mycobacterial DciA are essential for the interaction between DciA and the replicative helicase DnaB (35). Therefore, the DciA homologs in mycobacteria and the γ-proteobacteria *P. aeruginosa* and *V. cholerae* have diverged in the relative position of the DciA domain and the presence of N- or C-terminal sequence extensions. These sequence variations raise questions of whether there are functional consequences of these differences and if a broader view of the DciA protein family would reveal further diversification. To begin to address these open questions, we took a computational evolutionary approach and analyzed the phylogenetic distribution, domain architecture, and conservation of sequence properties amongst 26,789 DciA homologs. Our analysis revealed that most bacterial DciA proteins encode a single annotated domain, the eponymous DciA domain. However, we also identified multiple DciA homologs with novel domain architectures, which could provide clues to specialized functions or biology in those bacteria. Amongst the bacterial DciA single-domain proteins, there was lineage-specific variation in total protein length and positioning of the DciA domain. Despite this variation in sequence properties, AlphaFold structural predictions (37) identified a broadly conserved pattern of structural motifs where the DciA domain is connected to alpha-helical structures via an unstructured linker. Therefore, our analyses reveal conserved sequence-structural features of DciA homologs across bacterial phyla as well as lineage- and, sometimes, species-specific features, highlighting the need for expanded studies of this protein family.

## RESULTS

### DciA domain-containing proteins are predominantly found in bacteria with rare transfers to eukaryota

A protein homology search with *M tuberculosis* DciA (<u>AAK44227.1</u>) only identifies closely related homologs, predominantly in actinobacteria; similarly, homologs of *P. aeruginosa* DciA (<u>AAG07793.1</u>) are mostly proteobacterial. These data suggest that the DciA homologs across different phyla exhibit low conservation in their primary amino acid sequence. This divergence warranted a more comprehensive approach that used multiple DciA-domain containing proteins as starting points to retrieve the rich repertoire of DciA-like proteins from across the tree of life. Therefore, we analyzed the ∼27K InterPro entries for DciA domain-containing proteins (DUF721; Dna[CI] antecedent DciA; Pfam ID entry <u>PF05258</u>) (38), excluding metagenome data entries. We also added valuable metadata to this dataset, including protein accession numbers, taxID, species, and complete and collapsed lineages for each protein **(Table S1,** https://github.com/JRaviLab/dcia_evolution). Our initial characterization revealed that these ∼27K DciA-like proteins spanned the three kingdoms of life with 37 bacterial (with assigned lineages), 76 bacterial candidatus lineages (yet unassigned), 5 archaeal, 1 viral, and 6 eukaryotic lineages.

Next, we analyzed the phyletic distribution and domain architectures of all ∼27K DciA proteins using the MolEvolvR web application (39). This analysis highlighted that most (99.9%) proteins were found in the kingdom eubacteria (**Fig. 1A**). In addition, there were 26 DciA domain-containing proteins from eukaryota, archaea, and viral sequences (**Fig. 1A** **and Table S2)**. We performed BlastP homology searches with the 26 non-bacterial DciA domain-containing proteins (40) and found that over 95% of top hits with the highest confidence levels retrieved for 24 of these proteins were from bacterial genomes, suggesting that these DciA-domain containing proteins had been mistakenly attributed to archaeal, eukaryotic, or viral genomes **(Table S2)**. The two remaining non-bacterial DciA-like proteins occur in *Kingdonia uniflora* (KAF6150485.1), an endangered angiosperm with over-representation in DNA replication and repair genes (41), and the fungus *Hyaloscypha bicolor E* (<u>PMD57303.1</u>) **(Table S2, Fig. S1A)**. All close homologs of the *K. uniflora* DciA protein were in magnoliopsida, a class of flowering plants. The DciA protein from *H. bicolor E* predominantly retrieved (98/100) DciA-like proteins in fungal species. We also verified that these two eukaryotic DciA proteins were part of annotated ORFs in complete genomes, suggesting that they are truly eukaryotic proteins. Further, using AlphaFold structure prediction, we found that the DciA domains within the *K. uniflora* and *H. bicolor E* proteins resembled a truncated version of the *V. cholerae* DciA domain, solved previously using NMR (36) **(Fig. S1B)**. Each eukaryotic DciA domain is approximately half of the median length of *V. cholerae* DciA domain and is predicted to contain just the first alpha-helix and beta-sheet present in the *V. cholerae* DciA domain **(Fig. S1B)** (36). Our analysis using MolEvolvR revealed that the DciA domain-containing protein in *K. uniflora* also carries an annotated HMA (heavy metal associated) domain (Pfam: <u>PF00403</u>), overlapping with the DciA domain **(Fig. S1A,B)**. HMA domains are typically involved in heavy metal transport and detoxification in both eukaryotic and prokaryotic species (42-44), where heavy metal exposure can induce DNA damage (45-47). Although there were no additional Pfam domains annotated in the *H. bicolor E* DciA domain-containing protein, we identified two AN11006-like domains (PANTHER family: <u>PTHR42085</u>) of unknown function on either side of the DciA domain **(Table S2)**. Given that the DciA domain was previously considered to be an exclusively bacterial protein domain, the identification of predicted DciA structural domains in two eukaryotic species reveals the potential for much broader evolutionary distribution than previously appreciated, providing avenues for future structure-function studies.

**Figure 1.**
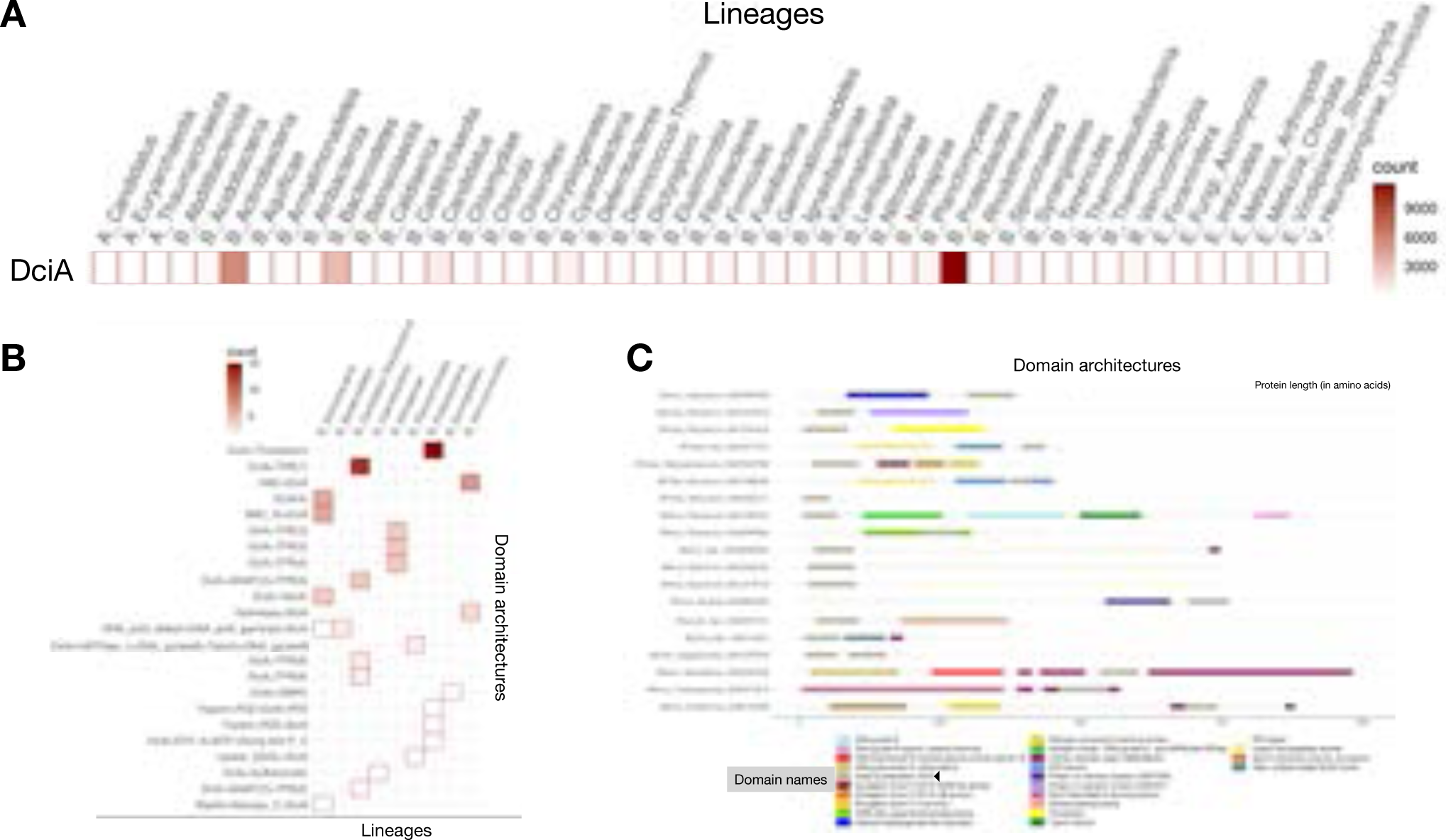
DciA protein phyletic spread and domain architectures. **A.** Phyletic spread of DciA domain containing proteins. The heatmap shows the abundance of DciA containing proteins across Bacterial, Archaeal, Viral, and Eukaryotic lineages. The color gradient indicates the number of homologs in a particular lineage. **B.** Phyletic spread of diverse DciA domain architectures (excluding DciA single-domain proteins) across bacterial lineages. The color gradient indicates the number of homologs with that domain architecture in a particular lineage. Domain architectures involving TPR repeats were combined based on the number of repeats. **C.** Key domain architectures of bacterial DciA homologs (including Pfam domains and MobiDBLite predicted IDRs). Representative proteins for each Pfam domain architecture within annotated bacterial lineages (no candidatus or metagenomes) were selected. Each representative protein is marked with the kingdom (B, bacteria), phylum (first 6 letters), Genus, and species (represented as ‘Gspecies’), and the NCBI protein accession number. The Pfam and MobiDB annotations for each domain prediction are shown in the legend. The arrow lengths represent the overall protein length. The characteristic DciA domain is indicated with a black arrow in the legend (grey domains).

### Bacterial DciA homolog domain architectures

#### DciA domains are mostly loners

The bacterial DciA proteins from InterPro fell into 37 bacterial lineages (**Fig. 1A**), plus many sequences annotated as Candidatus due to incomplete taxonomic classification **(Table S1,** https://github.com/JRaviLab/dcia_evolution). In line with sequencing and publication bias, proteobacterial, actinobacterial, and bacteroidetes genomes were over-represented in our homolog space (48-50) (**Fig. 1A**). Given the variation noted in the two eukaryotic DciA-domain containing proteins, we then proceeded to characterize the domain architecture of the ∼26K bacterial DciA homologs using MolEvolvR (39). We found that a stark majority (99.7%) of bacterial DciA proteins carried a lone DciA domain. In addition, we identified 38 variations where DciA homologs either contained multiple DciA domains or additional annotated domains (**Fig. 1B,C**). The domain architecture variations were often lineage-specific, as described below, indicating that they evolved later during speciation to adapt to lineage-specific biology.

#### Proteobacterial-specific variations in DciA domain architecture

Within proteobacteria, we identified four lineage-specific DciA domain architectures (**Fig. 1B**), including some domains present in other proteins associated with DNA replication. Specifically, 15 proteobacterial DciA homologs contained a thioredoxin-like domain (Pfam: Thioredoxin (TRX) <u>PF13462</u>; *e.g.,* ODT99650.1, *Rhodospirillales*) (**Fig. 1B,C**). TRX domains are present in a large class of redox-regulated proteins (51), and play roles in oxidative stress responses (52,53). In addition, *Cc*TRX1, an essential TRX domain protein, is upregulated during DNA replication initiation in the α-proteobacteria *Caulobacter crescentus* (54). Two other proteobacterial DciA homologs contain PDZ domains (Pfam: PF13180; e.g., KQZ00574.1, *Pseudolabrys*) (**Fig. 1B,C**), which are generally involved in protein-protein interactions (55,56) and could facilitate the interaction between these DciA proteins and other replication proteins. The PDZ domain-containing DciA homologs also carry trypsin-like peptidase domains (Pfam: PF13365) (**Fig. 1B,C**), which have recently been linked to DNA replication in humans, where the trypsin-like peptidase domain in the protein FAM111A is necessary for overcoming replication fork stalling (57). The DciA homolog in the α-proteobacteria *Micavibrio aeruginosavorus* (PZQ43964.1) encodes three translation elongation factor P domains (KOW-like, Pfam: PF0820, OB, Pfam: PF01132, C-terminal, Pfam: PF09285) (**Fig. 1B,C**).

#### Actinobacterial variations in DciA domain architectures

We identified four distinct lineage-specific domain architectures within actinobacteria, all of which involve domains associated with nucleotide sensing and DNA replication. Three Streptomyces DciA homologs contain a YspA domain (Pfam: <u>YAcAr/PF10686</u>; *e.g.,* SNC77843.1) in addition to the DciA domain (**Fig. 1B,C**). YspA domains typically have fusions to domains that sense and process nucleotide-derived ligands such as ADP-ribose (58). We found that the DciA protein in *Bifidobacterium callitrichos* (PST49340.1) contains the C-terminal domain of the DEAD-box RNA helicase family (Pfam: <u>PF00271</u>) (59), and the res subunit of the type III restriction enzyme, which encodes ATPase activity (Pfam: <u>PF04851</u>) (60) (**Fig. 1B,C**). In addition, five actinobacterial DciA homologs contain an N-terminal RecF/RecN/SMC domain (Pfam: <u>02463</u>; *e.g.,* OUE04448.1, *Clavibacter*) (**Fig. 1B,C**) common in the N-termini of structural maintenance of chromosome (SMC) proteins. The SMC domain typically includes an NTP-binding motif and SMC proteins are involved in chromosome partitioning, DNA recombination, and repair (61-65). In addition, six actinobacterial DciA homologs contain multiple DciA domains (e.g. OEJ23183.1, *Streptomyces*) (**Fig. 1B,C**).

In addition to actinobacteria-specific domain architectures, DciA proteins in *Micromonospora endolithica* (<u>RKN42798.1</u>) and bacteroidetes species *Rhodothermus marinus* (BBM73918.1, ACY49481.1) contain the γ/τ (Pfam: PF12169) and δ(Pfam: PF13177) subunit domains of DNA polymerase III (**Fig. 1B,C**), which make up part of the clamp loader complex in *E. coli* (66,67). The DNA pol III domains present in these DciA homologs also contain AAA+ ATPase domains, also found in DnaC/DnaI helicase loaders from *E. coli* and *B. subtilis* (29, 32, 33, 67).

#### Other DciA domain architectures involving domains associated with DNA replication

Thirty-six DciA homologs from nitrospinae and candidatus rokubacteria were annotated to contain tetratricopeptide repeat (TPR) domains (**Fig. 1B,C**). TPR domains are largely eukaryotic protein-protein interaction domains (68). For example, the TPR domain of the replication regulator Dia2 in eukaryotes is essential for the association of Dia2 with the replisome progression complex, which interacts with the MCM2-7 helicase at the replication fork (69). TPR proteins have also been identified in bacteria. For example, in the α-proteobacteria *Orientia tsutsugamushi*, two TPR proteins bind to the eukaryotic RNA helicase DDX3 to inhibit host cell translation (70). In candidatus rokubacteria, these TPR repeats are sometimes associated with an anaphase-promoting complex domain (Pfam: <u>PF12895</u>; *e.g.,* OLB41025.1), which along with TPR repeat domains, is present in eukaryotic cell cycle regulators (71,72).

The DciA homologs from two planctomycetes (KAF0244841.1, NUN49423.1) contain a Histidine kinase-, DNA gyrase B-, and HSP90-like ATPase domain (Pfam: <u>PF02518),</u> a DNA gyrase B domain (Pfam: PF00204), a toprim domain (DNA topoisomerase II Pfam: PF01751), and a DNA gyrase B C-terminus domain (Pfam: PF00986). A closer look at one of the planctomycetes DciA sequences (KAF0244841.1) shows that DciA and GyrB are annotated as a single fused protein. GyrB, part of the bacterial gyrase, is responsible for the negative supercoiling of dsDNA, which is essential during the opening of the DNA replication bubble (73).

#### Homology search with multiple starting points identifies additional DciA domain architectures

To ensure that we identified all sequenced DciA homologs and the entire repertoire of domain architectures in this protein family, we selected 94 unique lineage-domain-architecture pairs from 36 bacterial lineages and 27 total domain architectures to identify novel DciA homologs across the tree of life using MolEvolvR **(Fig. S2A)**. The only phylum without a representative from our initial analysis was atribacterota, which did not contain class-level assignments. In addition to novel domain architectures discussed above, the homology search identified 8 new domain architectures, including 11 actinobacterial homologs with a DciA dyad (*e.g.,* WP_165395910.1 from *Streptomyces*), several fusions with TPR repeats, and a DciA domain fused with an aminomethyltransferase domain (*Micrococcus sp. HSID17227*, WP_238693333.1) **(Fig. S2B)**. Together, our initial characterization along with these homologs captured the rich repertoire of DciA variation.

#### DciA single-domain homologs vary in protein length and position of the DciA domain

Despite having only one annotated domain, the DciA single-domain proteins varied considerably in length, ranging from 32 to 482 amino acids (**Fig. 2A**). In addition to total protein length, the distance of the DciA domain from the N- or C-terminus varied widely, where the DciA domain sometimes fell in the middle, at the N-terminus, or at the C-terminus of the protein (**Fig. 2A**). The median amino acid length of the annotated DciA domains in our dataset is 85aa **(Table S1)**, indicating that DciA homologs close to this protein length would only contain the DciA structural domain. An example of a DciA protein where the total protein length is similar to the size of the single DciA structural domain is the *Bacteroides fragilis* DciA homolog (<u>AUI45441.1</u>), which is 96aa long with the DciA domain annotated from amino acids 8-95. The AlphaFold structural prediction tool predicts that the entire *B. fragilis* DciA protein is comprised of the predicted DciA domain structure, consisting of one alpha-helix followed by a two beta-sheet motif, one kinked alpha-helix, and a third beta-sheet (34,36) (**Fig. 2B**). However, with a median protein length of 148aa (**Fig. 2A**), most DciA single-domain proteins are longer than the typical DciA domain length, suggesting that there may exist other functional sequences in DciA single-domain proteins that are not annotated as domains. For example, *V. cholerae* DciA is 157aa long, with the DciA domain positioned at the N-terminus (amino acids 12–90) followed by a 67aa C-terminal sequence extension. An NMR structure of the N-terminus of *V. cholerae* DciA confirms that the DciA domain in the N-terminus is sufficient to form the DciA domain structure (36). In addition to confirming the DciA domain structure in the N-terminus, AlphaFold prediction of the full-length *V. cholerae* DciA protein (<u>AAF95538.1</u>) depicts the C-terminus as an alpha-helix immediately following the DciA domain, connected to two terminal alpha-helices by an 11aa linker (**Fig. 2B**). Given that the C-terminus of *V. cholerae* DciA is required for its interaction with the replicative helicase *in vitro* (35), these data demonstrate not only that most DciA single-domain homologs are longer than the DciA domain alone, but also that the sequence extensions appended to the DciA domain can be physiologically relevant. The identification of DciA homologs with different domain architectures (**Fig. 1**), varying protein lengths (**Fig. 2A**), and different positioning of the DciA domain (**Fig. 2A**) suggest that some DciA homologs have evolved sequence properties with likely functional consequences for DciA activity.

**Figure 2.**
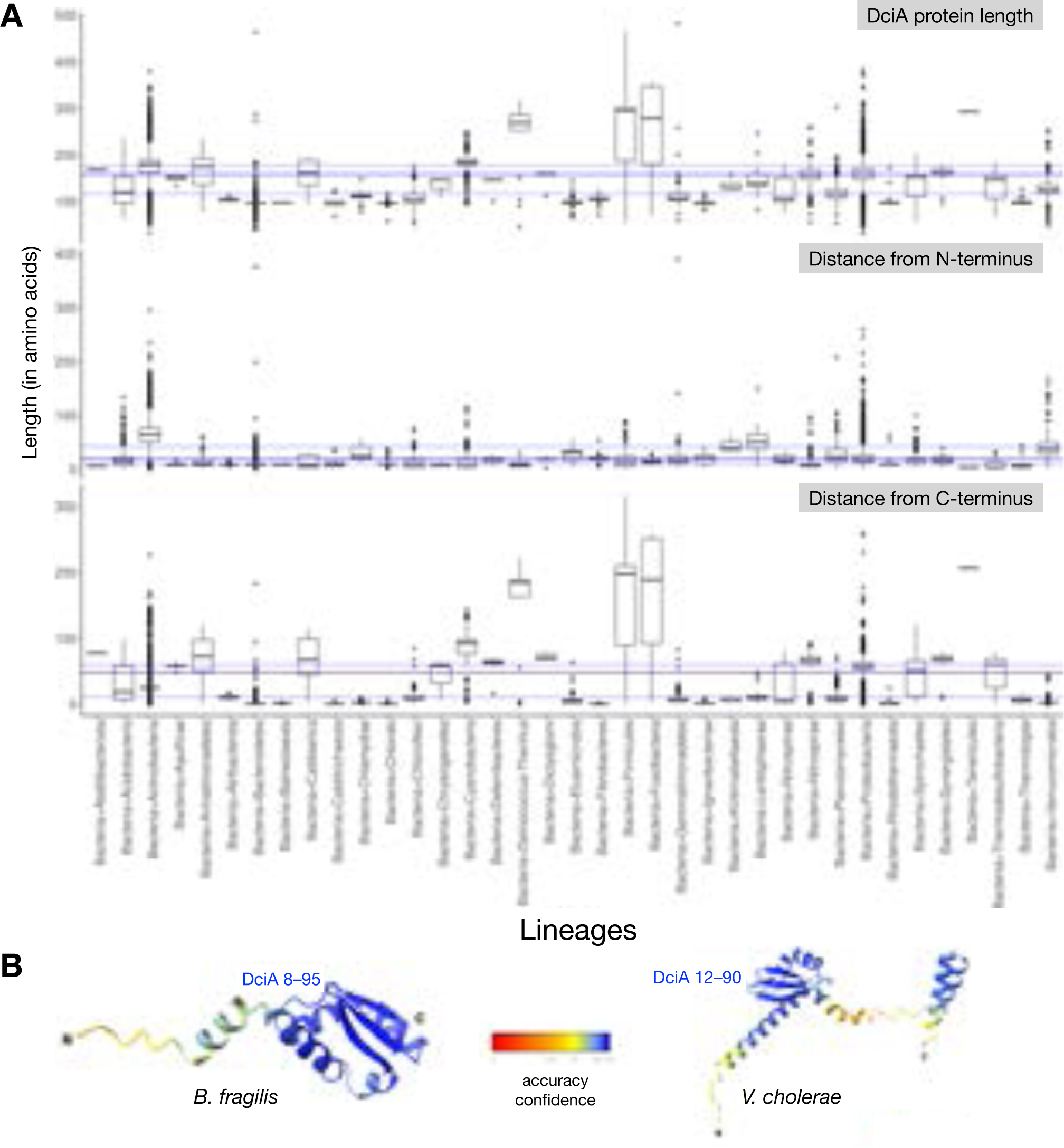
Length and DciA domain positioning in DciA single-domain proteins. **A.** Distributions of DciA single-domain protein length and the distances of the DciA domain from the N and C termini across bacterial lineages. Summary statistics for the single-domain DciA proteins were calculated for all bacterial lineages (no candidatus or metagenomes). The blue lines represent the 25th and 75th (light blue), and median/50th (dark blue) percentiles across lineages. **B.** AlphaFold structural prediction on the *B. fragilis* (<u>AUI45441.1</u>) and *V. cholerae* (<u>AAF95538.1</u>) DciA proteins visualized with ChimeraX. Protein is oriented left-to-right N-C termini. Color key indicates accuracy confidence (0-100).

#### Many DciA single-domain homologs are predicted to encode intrinsically disordered regions

Small-angle X-ray scattering, intrinsic disorder prediction tools, and molecular dynamic simulations have shown that the C-terminus of *V. cholerae* DciA that is required for DnaB binding *in vitro* contains an IDR (35). In addition, there is precedent for IDRs in other bacterial DNA replication proteins, including SSB (74-77), the replication restart helicase Rep (78,79), the helicase loaders DnaC and DnaI, and the replication initiation protein DnaA (35). To estimate the co-occurrence of IDRs with DciA domains, we used MobiDBLite (from within MolEvolvR (80)) to predict IDRs in each of the bacterial DciA homologs **(Table S1;** **Fig. 3A**). Out of ∼23K total bacterial non-candidatus DciA single-domain homologs, ∼8K proteins contained at least one IDR, and some (4,728) had multiple regions of disorder predicted. These IDRs were present in DciA homologs from 24 phyla (**Fig. 3A**). Analysis of the lengths of the IDRs in the DciA homologs from each phyla found that these IDRs ranged in length from 14aa to 213aa (**Fig. 3B**). *M. tuberculosis* DciA was among the single-domain proteins predicted to contain IDRs both N- and C-terminal to the DciA domain. Deletion of the N-terminal sequence of *M. tuberculosis* DciA that encodes the predicted IDR renders mycobacteria nonviable, demonstrating that this region predicted to form an IDR in DciA is essential for its cellular function (34).

**Figure 3.**
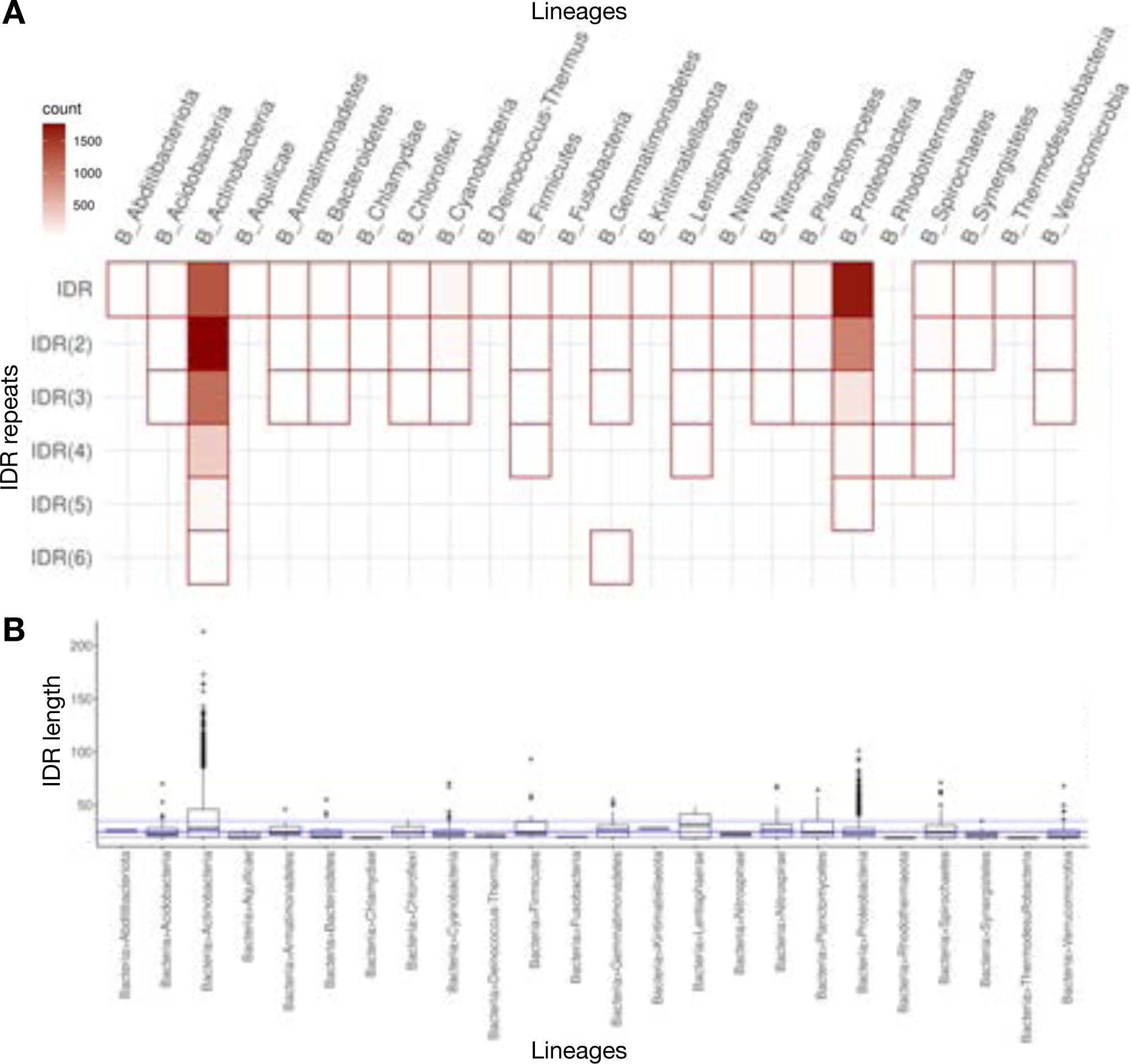
Disordered regions in DciA single-domain proteins. **A.** Phyletic spreads of Intrinsically disordered region (IDR)-containing DciA proteins. The heatmap shows the abundance of IDR-containing DciA proteins across bacterial lineages (no candidatus or metagenomes). ‘x’ in ‘IDR(x)’ indicates the number of IDRs predicted. The color gradient indicates the number of homologs in a particular lineage. **B.** Distribution of IDR length across bacterial lineages. Summary statistics for the single-domain DciA proteins with IDRs were calculated and plotted for each bacterial lineage. The blue lines represent the 25th and 75th (light blue), and median/50th (dark blue) percentiles across lineages.

#### DciA single-domain homologs can be separated into four groups based on protein length and the positioning of the DciA domain

The importance of the predicted IDR sequences for DciA activity in *M. tuberculosis* (34) and *V. cholerae* (35) along with the prevalence of predicted IDRs in DciA homologs suggest that IDRs in other DciA homologs may also be functionally relevant. In addition, we hypothesized that the N- and C-terminal sequence extensions in DciA single-domain proteins without predicted IDRs may also be functionally relevant. Since the minimum IDR sequence length in DciA homologs was 14aa (**Fig. 3B**), we reasoned that any N- or C-terminal sequence extension ≥14aa in a DciA protein could comprise a functionally relevant sequence. To identify DciA proteins with potentially relevant sequence extensions associated with the DciA domain, we binned the bacterial non-candidatus DciA single-domain homologs (23,309 total proteins) into four groups: i) Group 1 proteins with ≥14aa N-terminal to the DciA domain and <14aa C-terminal, ii) Group 2 proteins with <14aa N-terminal to the DciA domain and ≥14aa C-terminal, iii) Group 3 proteins with ≥14aa both N- and C-terminally to the DciA domain, and iv) Group 4 proteins with <14aa on both N- and C-termini (**Fig. 4 top** **panel, Table S1)**.

**Figure 4.**
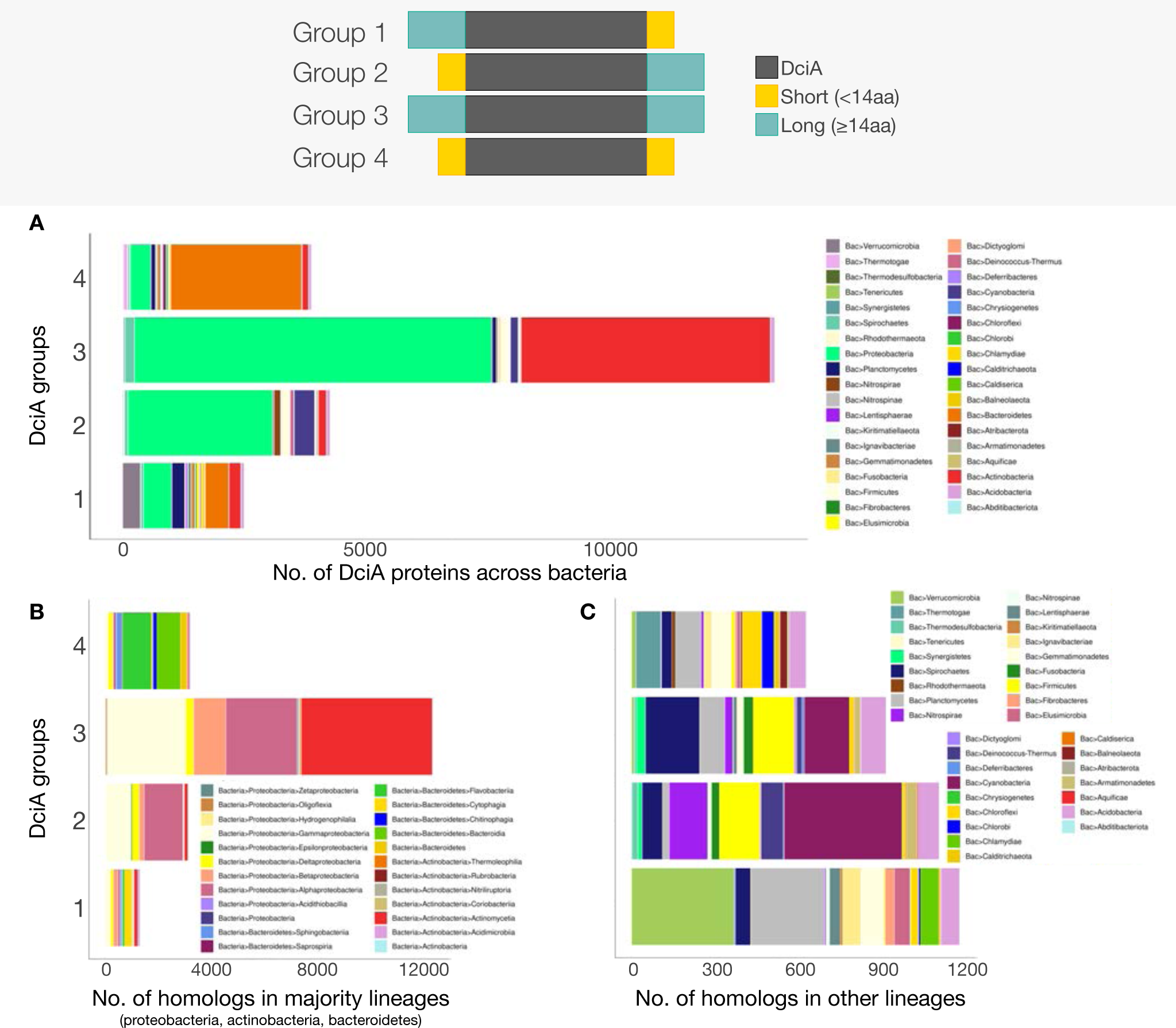
Grouping of DciA single-domain proteins and their phyletic spread. The top panel shows the four Groups of bacterial DciA single-domain proteins binned based on the lengths of flanking N- and C-terminal extensions of DciA domains: : i) Group 1 proteins with ≥14aa N-terminal to the DciA domain and <14aa C-terminal, ii) Group 2 proteins with <14aa N-terminal to the DciA domain and ≥14aa C-terminal, iii) Group 3 proteins with ≥14aa both N- and C-terminally to the DciA domain, and iv) Group 4 proteins with <14aa on both N- and C-termini. Stacked barplots of 4 groups of DciA single-domain proteins are plotted across **A.** all bacterial lineages, with a focus on **B.** predominant lineages further resolved into sub-phyla (proteobacteria, actinobacteria, and bacteroidetes), and **C.** other lineages. The number of homologs in each group is are further characterized based on their lineage-wise distribution. No bacterial candidatus or metagenomes are displayed.

Most (∼90%) actinobacterial DciA homologs fell into Group 3, with sequences on both sides of the DciA domain, although there were examples of actinobacterial DciA proteins in each of the other three groups as well (**Fig. 4A**). When we separated actinobacteria by class, we found that actinomycetia and coriobacteria DciA proteins were mostly in Group 3, whereas acidimicrobia, nitriliruptoria, and thermoleophilia DciA proteins were mostly in Group 1 (**Fig. 4B**). Group 3 also contained several (∼65%) proteobacterial DciA homologs, however, there were also 2,958 proteobacterial DciA proteins in Group 2, as well as representatives in Groups 1 and 4 (**Fig. 4A**). When we separated proteobacteria into individual classes, homologs from α-proteobacteria, β-proteobacteria, δ-proteobacteria, γ-proteobacteria, oligoflexia, and ζ-proteobacteria fell mostly into Group 3 (**Fig. 4B**). In contrast, DciA homologs in ε-proteobacteria and hydrogenophila mostly fell into Group 2. In contrast to both actinobacteria and proteobacteria, bacteroidetes DciA homologs fell almost exclusively into Groups 1 and 4, with the majority (∼84%) belonging to Group 4 (**Fig. 4A**). When we separated bacteroidetes into classes, we found that only the cytophagia bacteria contained mostly Group 1 homologs, while the rest of bacteroidetes classes contained mostly Group 4 homologs, indicating lineage and class-specific selection of DciA homolog sequence structures (**Fig. 4B**).

To better visualize the distribution of DciA single-domain proteins from phyla with smaller numbers of representative sequences, we removed the proteobacterial, actinobacterial, or bacteroidetes sequences and plotted the distribution of the remaining DciA single-domain homologs in Groups 1–4 (**Fig. 4C**). Some lineages had almost all DciA homologs classed within a single Group, indicating the evolution and conservation of a particular DciA sequence organization for that lineage. For example, 94% of verrucomicrobia and 96% of chlamydiae DciA homologs fell into Group 1. Other lineages had DciA homologs more broadly distributed across Groups, although there were still lineage-specific biases towards single Groups. Greater than 65% elusimicrobia, lentisphaerae, ignavibacteriae, and fibrobacteres DciA proteins were classed into Group 1, >65% of cyanobacteria, nitrospirae, deinococcus-thermus, and thermodesulfovibrio DciA homologs were classed in Group 2, and >65% of synergistetes DciA homologs are in Group 3. Planctomycetes exhibited differentiation of DciA homolog grouping at the Class level. Planctomycetes class planctomycetia homologs were mostly in Group 1, while planctomycetes phycisphaerae homologs were split evenly between Groups 1 and 3 (**Fig. 4C**, **Table S1**). Firmicute and fusobacteria DciA homologs were represented equally in Groups 2 and 3, with a small number of representatives in groups 1 and 4. Alternatively, gemmatimonadetes DciA homologs were predominantly split between Groups 1 and 4, and nitrospinae homologs were split between groups 1 and 2.

In addition to bacteroidetes, other lineages that were predominantly in Group 4 included chloroflexi (56% of DciA homologs), rhodothermaeota (81%), calditrichaeota (74%), chlorobi (86%), and thermotogae (91%) (**Fig. 4C**). Group 4 proteins only encode the DciA structural domain, indicating that the DciA domain is sufficient for DciA activity in these bacteria, although this has yet to be experimentally tested. DciA homologs from acidobacteria, spirochaetes, and armatimonadetes did not follow any apparent trends in DciA homolog classification, and kiritimatiellaeota, aquificae, chrysiogenetes, abditibacteriota, tenericutes, balneolaeota, caldiserica, dictyoglomi, atribacterota, and deferribacteres had <20 sequences in our dataset. The lineage and class-specific trends observed in the four different DciA groupings suggest differentiation of DciA proteins in a largely lineage-specific manner. The sequences N- and/or C-terminal to the DciA domain in Groups 1–4 could have functional consequences for the mechanisms of DciA proteins in different bacterial lineages.

#### Structure predictions of DciA single-domain homologs reveal conserved structural motifs within and outside of the DciA domain

To understand the impact of sequence extensions on protein structure in DciA single-domain homologs, we used AlphaFold structure prediction to model representative DciA proteins from Groups 1–4 (**Fig. 5**). For Groups 1–3, we also compared the structures of homologs with and without IDRs predicted in the sequence extensions. Regardless of Group designation, all DciA domains were predicted to fold into a structure similar to that previously predicted for *M. tuberculosis* DciA (34) and experimentally validated for *V. cholerae* DciA (36) (**Fig. 5**).

**Figure 5.**
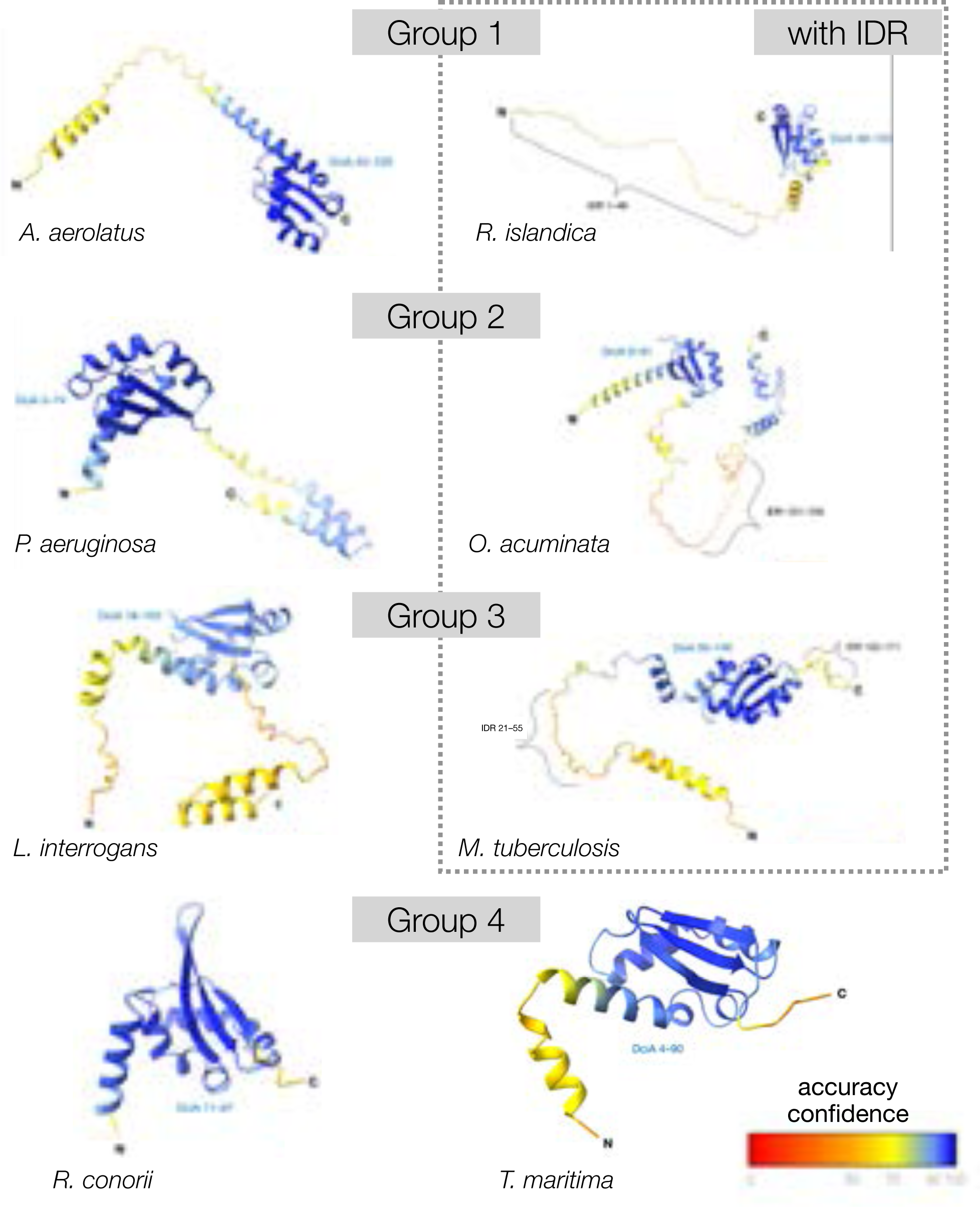
Structure prediction of representative DciA homologs by groups (with and without IDR). AlphaFold representatives from **A.** Group 1 DciA proteins from *A. aerolatus* (left; <u>GEO04242.1</u>) and *R. islandica* (right; <u>KLU02380.1</u>), **B.** Group 2 DciA proteins from *P. aeruginosa* (left; <u>AAG07793.1</u>) and *O. acuminata* (right; <u>AFY85399.1</u>), **C.** Group 3 DciA proteins from *L. interrogans*(left; <u>AAN47203.1</u>) and *M. tuberculosis* (right; <u>BAL63824.1</u>), and Group 4 DciA proteins *T. maritima* (left; <u>AAD35914.1</u>) and *R. conorii* (right; <u>AAL03818.1</u>). Models visualized with ChimeraX. Proteins are oriented left-to-right N-C termini, termini marked on each structure. DciA domains annotated in blue text, IDRs annotated with black bracket with all corresponding amino acid annotations. Key indicates accuracy confidence (0-100). See *Methods* for accession codes of each model deposited in ModelArchive.

For a Group 1 DciA from planctomycetes (<u>KLU02380.1</u>, *Rhodopirellula islandica*) that is predicted to have a 48aa IDR N-terminal to the DciA domain, AlphaFold predicts that the N-terminal sequence extension does not form any structured helices or beta sheets, supporting that this region could be intrinsically disordered (**Fig. 5**). In contrast, the structural prediction of a Group 1 DciA single-domain protein from bacteroidetes (<u>GEO04242.1</u>, *Adhaeribacter aerolatus*) with no predicted IDR displays the C-terminal sequence extension ending with a single alpha-helix, connected to the DciA protein with an unstructured linker (**Fig. 5**).

A Group 2 DciA homolog from cyanobacteria (<u>AFY85399.1</u>, *Oscillatoria acuminata*) that is predicted to contain a C-terminal IDR exhibited the classic DciA domain folds followed by one alpha helix connected to two more C-terminal helices via a 24aa linker, which coincided with the predicted IDR (**Fig. 5**). The Group 2 DciA homolog from *P. aeruginosa* (<u>AAG07793.1</u>, γ-proteobacteria) with no predicted IDR, contains the DciA domain folds connected to two C-terminal helices via a 16aa linker (**Fig. 5**), which is reminiscent of the structure for O. *acuminata* DciA,.

The Group 3 DciA protein from the actinobacteria *M. tuberculosis* (<u>BAL63824.1</u>) consists of a single 18aa alpha helix in its N-terminus that is connected to the DciA domain by a 34aa linker region, which is predicted to contain an IDR (**Fig. 5**). The C-terminus of *M. tuberculosis* DciA is also predicted to contain a 19aa IDR. A group 3 protein with no predicted IDR from *Leptospira interrogans* (spirochaete (<u>AAN47203.1</u>) contains sequence extensions in both its N- and C-terminus (**Fig. 5**), with the same double alpha helix fold in its C-terminus as observed in the group 2 proteins *P. aeruginosa* and *O. acuminata.* The DciA domain is also connected to the two-alpha helical domain by a 20aa unstructured linker region.

Similar to the structure predicted for *B. fragilis* DciA (**Fig. 2B**), other Group 4 DciA homologs from thermotogae (*Thermotoga maritima*, <u>AAD35914.1</u>) and α-proteobacteria (*Rickettsia conorii*, <u>AAL03818.1</u>) only contain the folds previously reported for the DciA domain in *V. cholerae* (36) (**Fig. 5**). These data further support that the DciA structural domain alone is sufficient for DciA activity in Group 4 bacteria.

Visualization of these representative DciA structures reveals that when unannotated sequences are appended to the DciA domain (Groups 1–3), they tend to form an unstructured linker region connected to 1–2 alpha-helices at the termini, regardless of whether the sequences are positioned N- or C-terminal to the DciA domain. In some Group 1–3 DciA homologs, the linker is predicted to comprise an IDR. Notably, *V. cholerae* DciA has an experimentally verified C-terminal IDR (35), although this was not predicted by MobiDBLite **(Table S1)**. Circular dichroism and secondary structure prediction tools of *V. cholerae* DciA predict that the C-terminal IDR can transiently form two alpha-helices (35) (**Fig. 2B**). These helices occur at a similar position in *P. aeruginosa* DciA as well (**Fig. 5**). Therefore, it is possible that the unstructured linker motifs in *P. aeruginosa* and other Group 1–3 DciA homologs comprise IDRs not predicted by MobiDBLite due to the alpha-helices that can form in the termini. The conservation of the predicted structures of DciA homologs across bacterial lineages, where the DciA domain is connected to alpha-helical structures via an unstructured linker, suggests that these structural motifs are important for DciA activity in bacteria that encode Group 1–3 homologs.

#### DciA evolution across the tree of life

Our sequence-structure analyses thus far revealed lineage-specific signatures in DciA domain organization in addition to the widely prevalent DciA single-domain proteins. The natural next, and final, question is how did these different DciA proteins evolve — are there species/lineage-specific, domain architecture, or group-specific migration patterns? To address these questions, we used all DciA proteins from **Fig. 1** to generate a phylogenetic tree (**Fig. 6**). Most strikingly, we observed that DciA homologs clustered by lineages (**Fig. 6A**). The three largest lineages, actinobacteria, bacteroidetes, and proteobacteria, are labeled based on their dominant membership (**Fig. 6A**). We noted two main proteobacterial clades (left and bottom), likely due to class-wise grouping. We, therefore, further resolved the tree by the predominant bacterial classes from these three phyla (**Fig. 6B**). The class-resolved tree explains the distinct proteobacterial clusters observed in the phylum-based tree (**Fig. 6A**), wherein α- proteobacteria and β/γ-proteobacteria form distinct clusters (**Fig. 6B**). This migration pattern of the α-proteobacterial homologs suggests that the DciA proteins in this lineage have diverged evolutionarily from the rest of the proteobacterial lineages. The phylogenetic analysis indicates that variations in DciA protein domain architectures, protein lengths, and DciA domain positioning likely occurred after lineage-specific divergence of bacterial classes. Overall, the DciA phylogenetic tree delineates the evolution of this critical panbacterial protein across all major lineages.

**Figure 6.**
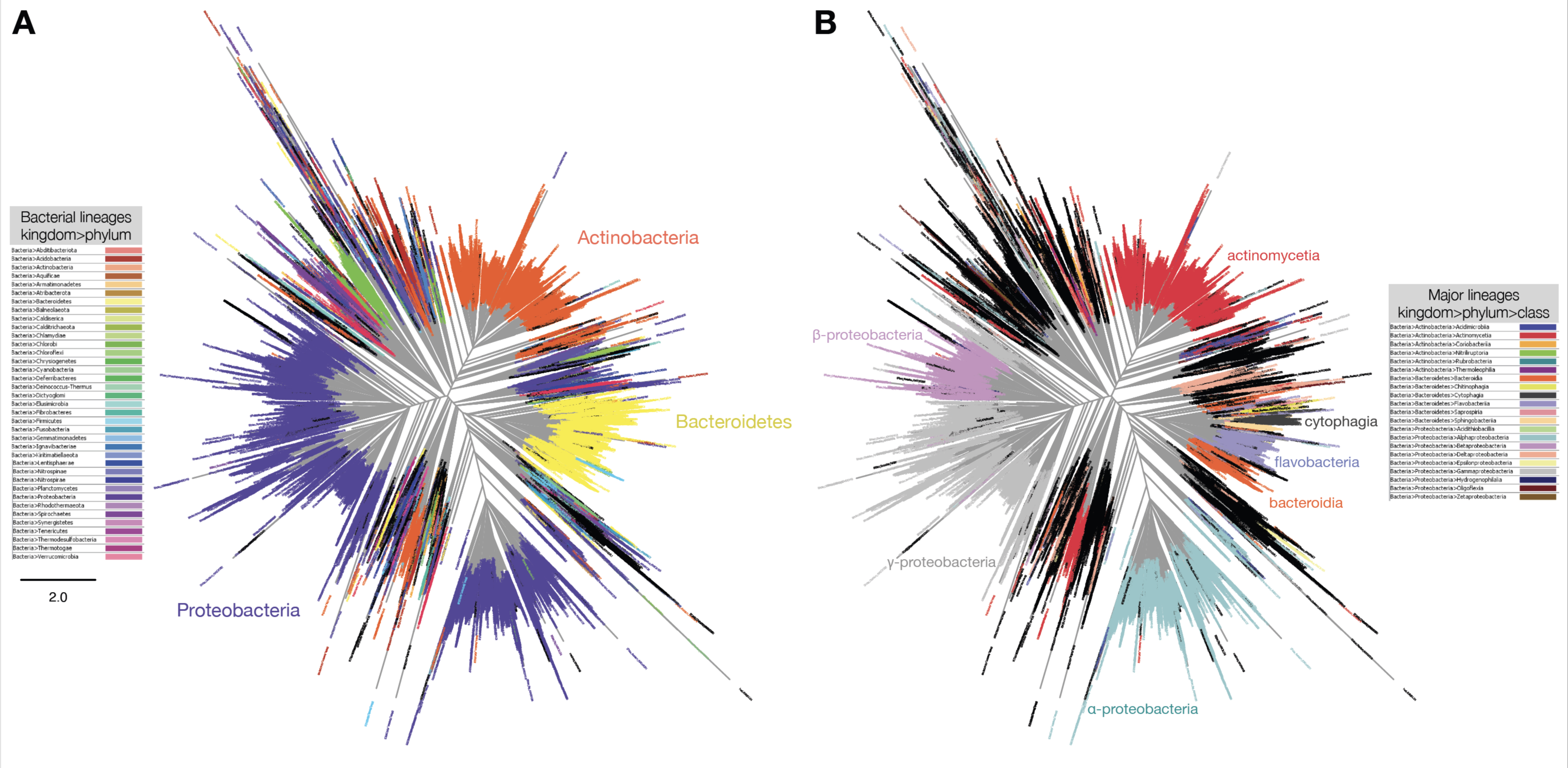
DciA evolution across the tree of life. All DciA domain-containing proteins from Figure 1 were used to reconstruct the DciA phylogenetic tree. Kalign3 (87) was used for multiple sequence alignment, and FastTree (88) was used to generate the tree. The resulting tree was visualized using FigTree (http://tree.bio.ed.ac.uk/software/figtree/). The tree was colored by major lineages (left: bacterial phyla; right: key bacterial classes) with the remaining DciA proteins in black. The three predominant bacterial lineages, proteobacteria, actinobacteria, and bacteroidetes, are marked on the phylum-based tree **(A)** next to their corresponding largest clusters, and the major bacterial classes are marked next to their largest clusters in the class-based tree **(B)**.

## DISCUSSION

The recent discovery of DciA as a predicted helicase operator in bacteria (28,34,36) has begun to shed light on a long-standing open question of how the majority of bacteria facilitate helicase activity during DNA replication in the absence of the ATPase helicase loaders expressed by *E. coli* and *B. subtilis*. The wide distribution of DciA in diverse bacterial phyla indicates that these proteins likely represent the predominant paradigm for helicase operation, despite not being conserved in *E. coli* and *B. subtilis,* the organisms typically used as a model for bacterial replication. DciA proteins are defined by the presence of the DciA domain (28). Prior phylogenetic analysis indicates that *dnaC*and *dnaI* homologs were acquired through evolution at the expense of *dciA* (named for *dna[CI]* antecedent) (28,30,31) suggesting that DciA and DnaC/DnaI perform a common function. In addition, it has been shown that DciA interacts with the replicative helicase and is required for DNA replication and viability in the few organisms it has been studied in (28, 34–36). However, the mechanism by which DciA mediates replication initiation is still unknown.

Our comprehensive evolutionary study of DciA proteins has revealed both lineage-specific and conserved features amongst homologs. We find that most homologs are DciA single-domain proteins in bacteria, with rare instances of additional domains in DciA homologs, many of which have known roles in DNA replication and repair **(Fig. 1, S2)**. These additional domain architectures were predominantly phyla-, and sometimes species-, specific, suggesting that they have been acquired and maintained to facilitate lineage-specific requirements during DNA replication. Further study of these DciA domain architectures could shed light on varying mechanisms of the regulation of bacterial replication, or other roles for DciA in the cell. Similarly, we identified two eukaryotic proteins that encode partial DciA domains **(Fig. S1)**, raising the question of how this domain would function in eukaryotes in the absence of its bacterial replicative helicase binding partner. The eukaryotic DciA-domain containing proteins also harbored additional domains without known connections to DNA replication, possibly suggesting that the DciA domain has been co-opted for other purposes in these organisms.

Even though most bacterial DciA homologs were single-domain proteins, they exhibited a wide variety in sequence lengths and positioning of the DciA domain (**Fig 2**, **Table S1)**. When we grouped the DciA homologs based on the position of the DciA domain and the presence of N- and C-terminal extensions, we identified lineage-specific trends in these sequence features (**Fig. 4**), suggesting that these variations mostly evolved following speciation and highlight how the regulation of helicase activity during DNA replication initiation could differ between phyla. For example, most actinobacterial and proteobacterial DciA single-domain homologs fell into Groups 2 and 3, which harbored sequence extensions either N-terminally or on both sides of the DciA domain, while most bacteroidetes DciA homologs were classed in Group 4 encoding only the DciA structural domain. The sequence extensions in actinobacteria *M. tuberculosis* and proteobacteria *V. cholerae* DciA have been shown to be essential for DciA activity *in vivo* or *in vitro*, respectively (34,35). The absence of these sequences in most bacteroidetes homologs suggests differing requirements for DciA activity in different bacteria.

Despite the lineage-specific grouping of DciA homologs based on sequence lengths and positioning of the DciA domain, AlphaFold structural prediction of representative DciA homologs from each Group revealed common structural patterns (**Fig. 5**). The DciA domain structure (36) was conserved across DciA homologs, and has previously been noted to resemble the structure of the N-terminus of DnaA (34–36, 81). The N-terminus of DnaA is critical for the interaction with the DnaB replicative helicase and other regulators (82,83), however, the role of the DciA domain in the interaction with DnaB has yet to be established. A tryptophan residue conserved in the DciA domains of many DciA homologs is positioned similarly to a phenylalanine residue in the DnaA N-terminus that has been predicted to have a key role in making contacts between DnaA and its interacting partners, including DnaB (34,84). Mutation of the tryptophan in the DciA domain of *M. tuberculosis* DciA results in slow growth and decreased DNA replication (34). This supports that the tryptophan within the DciA domain plays a key role for DciA function *in vivo* in mycobacteria. However, not all DciA homologs encode this tryptophan residue within their DciA domain, a key example being *V. cholerae* DciA, which has an isoleucine at this position (35). Therefore, even the defining DciA domain feature of DciA proteins exhibits some variation in different bacteria that may reflect differences in mechanism of action or mode of interaction with the replicative helicase.

In addition to conservation of the DciA domain structure, when sequence extensions were associated with the DciA domain, they tended to form unstructured linkers terminating in 1–2 alpha-helices (**Fig. 5**). These linker and alpha-helical structures were identified either C- or N-terminal to the DciA domain, depending on the homolog, where the role of this positioning in terms of function is still unknown. At least one third of the DciA single-domain homologs were also predicted to contain IDRs **(Fig. 3, 5)**. The IDR C-terminal to the DciA domain in the *V. cholerae* homolog is required for its interaction with the DnaB helicase (35). IDRs in other replication proteins, including SSB and Rep, also play key roles in facilitating protein-protein interactions within the replisome (74-79), indicating a common theme and function of these domains during bacterial DNA replication. However, if the IDR in the unstructured linker sequence is required for the interaction between DciA and the replicative helicase, this would imply that Group 4 DciA homologs, which only encode the DciA structural domain and no linker domains, employ other sequences or mechanisms to interact with DnaB. Thus far, no DciA homologs in Group 4 have been studied either genetically or biochemically and so how the DciA domain functions on its own remains a mystery.

Our computational evolutionary analyses have enabled an in-depth delineation of the evolution of DciA across the tree of life and elucidated the variation in DciA domain architectures, co-occurrences with IDRs, and the many flavors of bacterial DciA sequence-structural features. Despite this deep exploration, many unknowns still remain regarding DciA proteins and bacterial DNA replication. Our results highlight the complexities and diversity that have evolved in the fundamental process of DNA replication, where no single species of bacteria will be able to represent a central dogma that holds true throughout the kingdom. These studies provide a framework for researchers to consider the evolutionary variation while dissecting the mechanistic basis for helicase operation in bacteria.

## METHODS

### Query selection

We started with ∼27K DciA carrying proteins from Pfam (UniProt sequences from the InterPro database (38)). We added relevant metadata to each of these homologs, including corresponding NCBI protein accession numbers, protein length, taxID, species, and lineages. We used MolEvolvR (39) and the underlying InterProScan (85) to explore the domain architectures, cellular localizations, and disorder predictions for all ∼27K DciA proteins. The resulting domain architectures (from MolEvolvR) were also appended to each homolog’s metadata. All these data are available in **Table S1**, and on GitHub (https://github.com/JRaviLab/dcia_evolution).

### Representative DciA proteins

To select representatives for subsequent analyses, we picked DciA carrying proteins from each lineage–domain-architecture combination. We excluded proteins that were not bacterial, and those without an assigned lineage, (*e.g.,* candidatus and uncultured bacteria). The remaining sequences were then grouped by the short lineage column (superkingdom>phylum>class) and the Pfam domain architecture column. These groups were then reverse sorted by sequence completeness and one representative protein in each group was selected. From this selection process we found 94 representative proteins from 72 bacterial lineages with 23 unique domain architectures.

### Protein homology search

We performed homology searches of the 26 non-bacterial DciA proteins using the NCBI BLASTP (40). When examining the results, we excluded any non-bacterial query protein (from the 27) if they retrieved <5% non-bacteria in their top 100 hits (or 23 hits in the case of <u>RLI67678.1</u>) (≥95% were bacterial).

### Analysis using MolEvolvR

#### Domain architectures

We used MolEvolvR (39) to determine and characterize all DciA-containing proteins from InterPro and their homologs. We first downloaded the sequences of all DciA-containing proteins identified by InterPro from UniProt. The domain architectures were identified using InterProscan (85) and analyzed by lineage, quantified with heatmaps, and visualized by unique combinations of Pfam domains. During this analysis, we renamed MobiDBLite predictions to “Intrinsically disordered regions” to be consistent with previous literature. Also, DciA homologs with TPR fusions were condensed by the number of TPR repeats for clarity, *e.g.,* TPR+TPR+TPR was condensed to TPR(3). We then aggregated relevant species and protein annotation metadata from the NCBI into our combined dataset. We selected 94 representatives to include each combination of bacterial lineage (excluding atribacterota, which did not contain class-level assignments) and Pfam domain architecture as queries for MolEvolvR homology search.

#### Homology search

These 94 proteins were submitted to MolEvolvR to identify homologs in the NCBI RefSeq non-redundant proteins database (86). The domain architectures and sequence-structure motif predictions, as well as lineage and protein-related metadata, were determined using MolEvolvR for each of the homologs (as described above) for downstream analyses. The resulting data were summarized and visualized from within MolEvolvR and using custom R scripts.

#### Data availability

All our data, analyses, and visualizations summarizing the DciA homologs across the bacterial kingdom, along with their domain architectures and phyletic spreads, are available at https://github.com/JRaviLab/dcia_evolution. Detailed legends for our data tables, structure predictions, and sequences used for tree generation are also available in our GitHub repository.

### Phylogenetic Tree

Fasta sequences of all DciA-containing proteins from InterPro (∼27k) were obtained from the UniProtKB. These sequences were aligned using kalign3 (87) and a phylogenetic tree was constructed using FastTree (88). The resulting tree was visualized with FigTree (http://tree.bio.ed.ac.uk/software/figtree/) and color-coded by lineage.

### Alphafold Structure Prediction

We used the AlphaFold structural prediction <u>Colab notebook</u> (37,89) via the ChimeraX 1.4 daily build software (90) downloaded 2022-02-07 for all protein models. Visualization and analyses of models performed with UCSF ChimeraX, developed by the Resource for Biocomputing, Visualization, and Informatics at the University of California, San Francisco (90). All AlphaFold structures have been deposited via ModelArchive (http://modelarchive.org/), and are available with the following accession codes (follow the link and enter the temporary supplementary access code when prompted): <u>ma-q8sq3</u> (*K. uniflora* – temporary supplemental access code: xbO9Q6QacF), <u>ma-vyetl</u> *(H. bicolor E –* temporary supplemental access code: u1N27DoUat), <u>ma-z6rsv</u> *(B. fragilis* – temporary supplemental access code: 16QiFKRCzw), <u>ma-1hvpi</u> *(V*. *cholerae* – temporary supplemental access code: UWLLwXhQa4), <u>ma-tk4v8</u> *(R. islandica* – temporary supplemental access code: ljFtsjb0cW), <u>ma-9cnra</u> *(A. aerolatus* – temporary supplemental access code: uyu5HEqHrF), <u>ma-v2jc1</u> *(O. acuminata –* temporary supplemental access code: yMKAkPTopL), <u>ma-ibgex</u> *(P. aeruginosa –* temporary supplemental access code: HcHuzHbF4s), <u>ma-2ovw9</u> *(M. tuberculosis –* temporary supplementary access code: kUR6yS2EXW), <u>ma-n7tbg</u> *(L. interrogans –* temporary supplementary access code: R02X0DN45Q), <u>ma-02qnb</u> *(T. maritima* – temporary supplemental access code: LvLPofxN1e), <u>ma-3ee0e</u> *(R. conorii –* temporary supplementary access code: GBBQZBO9LD). Public DOIs (currently pending) will become available upon publication. The structures are also available in our GitHub repository (PDB format; under ‘model_structures’): https://github.com/JRaviLab/dcia_evolution.

## Supporting information

Supplemental Information Combined

## ACKNOWLEDGEMENTS

We are very grateful to the Midwest Microbial Pathogenesis Conference (MMPC) 2021 organizers for providing HCB a travel award and opportunity to present the DciA story. This interactive venue enabled the start of this collaboration between JR, JTB and CLS, HCB.

## DATA AVAILABILITY AND REUSE

All the data, analyses, and visualizations are available in our GitHub repository, https://github.com/JRaviLab/dcia_evolution. Text, figures, and data are licensed under Creative Commons Attribution CC BY 4.0.

