## Supplemental Information Combined for "DciA helicase operators exhibit diversity across bacterial phyla"

### Supplementary Figures

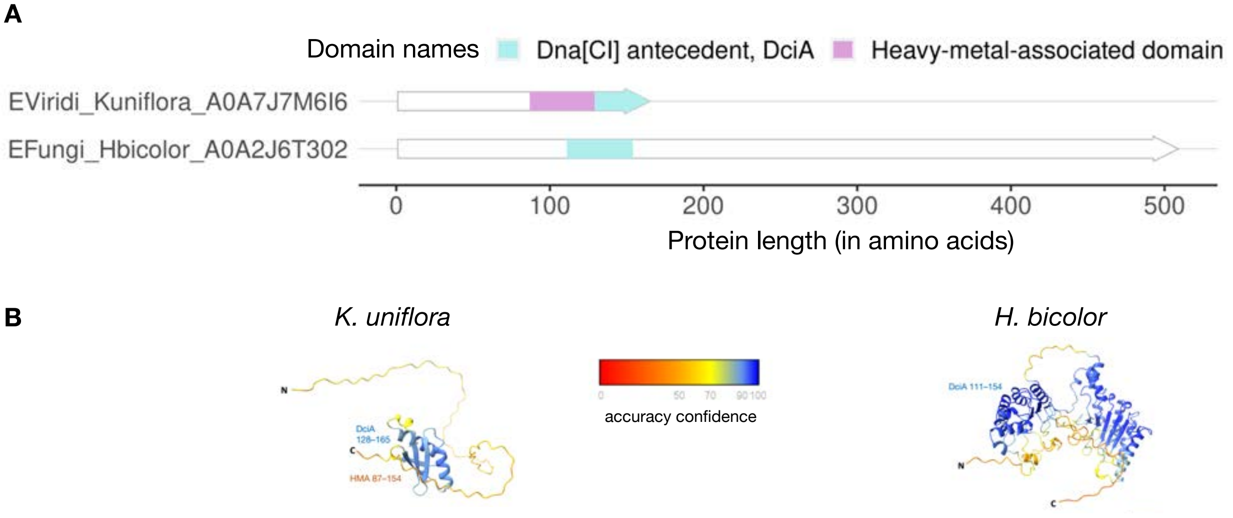


**Figure S1. Novel eukaryotic DciA domain architectures. A.** Pfam domain architecture visualization using MolEvolvR**.** Each DciA protein is marked with the kingdom (B, bacteria), phylum (first 6 letters), Genus, and species (represented as 'Gspecies'), and the NCBI protein accession number. The Pfam annotation for each domain prediction is shown in the legend (top). The arrow lengths represent the overall protein length. **B.** AlphaFold structure predictions for the DciA homologs in *K. uniflora* (left; KAF6150485.1) and *H. bicolor E* (right; PMD57303.1) proteins, visualized using ChimeraX. Proteins are oriented left-to-right N–C termini. Key indicates accuracy confidence (0-100). Domain predictions (from Pfam, using MolEvolvR) are marked on the structures.

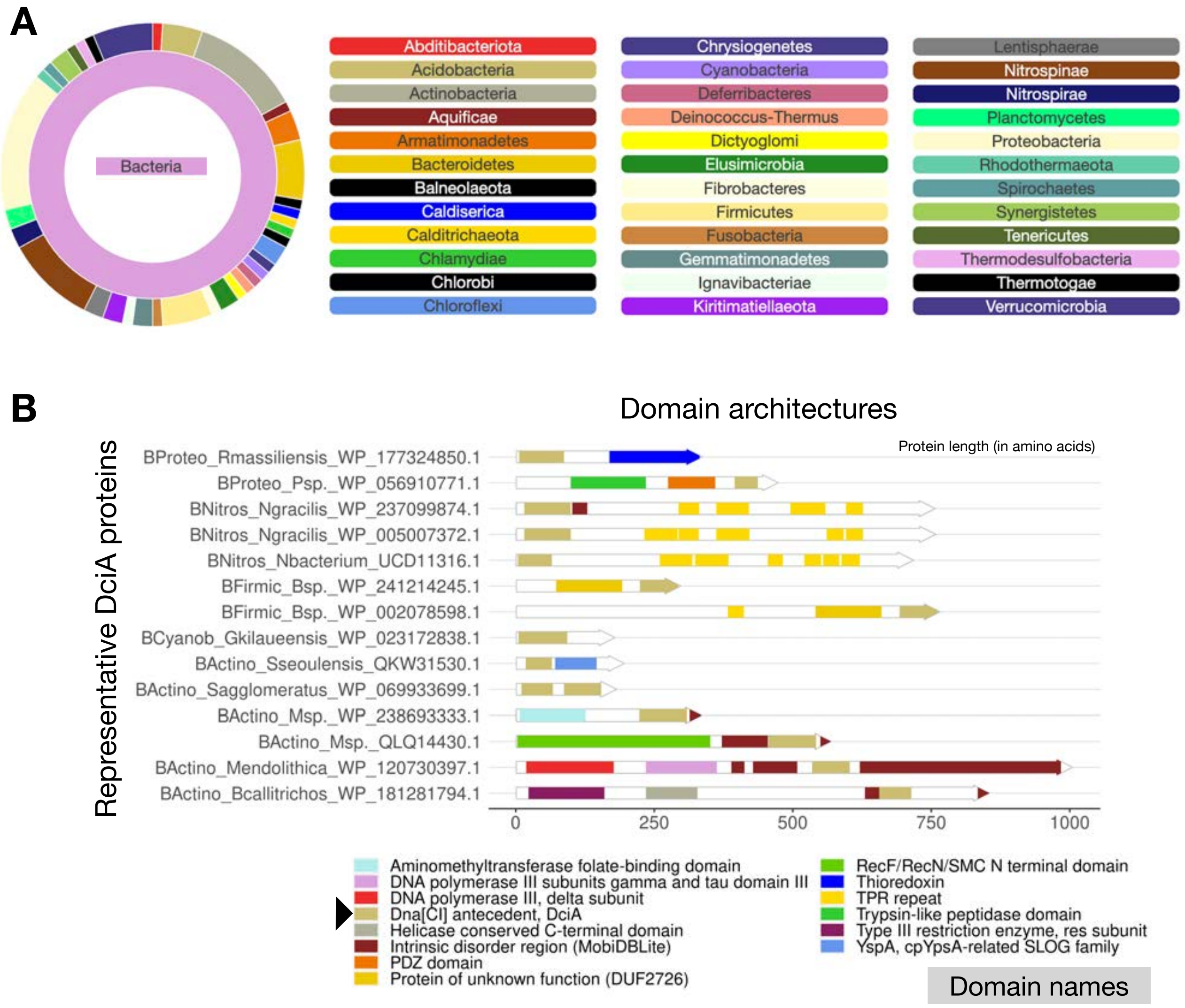


**Figure S2. Searching for DciA domain-containing proteins across the bacterial kingdom using MolEvolvR. A.** Bacterial lineages of representative DciA domain-containing proteins, serving as starting points used for the comprehensive homology search. The representative 94 DciA proteins include 23 domain architectures and 72 lineages (kingdom>phylum>class). The sunburst plot shows the fractional distribution of representative DciA across bacterial lineages (no metagenomes, or candidatus/unassigned bacterial lineages were included). The inner ring corresponds to the kingdom and the outer ring represents the distribution of phyla. **B.** Representative domain architectures of DciA domain-containing homologs from MolEvolvR (Pfam and MobiDBLite). Representative homologs for each Pfam–MobiDBLite domain architecture combination were selected. Each representative protein is marked with the kingdom (B, bacteria), phylum (first 6 letters), Genus, and species (represented as 'Gspecies'), and the NCBI protein accession number. The Pfam and MobiDBLite annotations for each domain prediction are shown in the legend (bottom). The arrow lengths represent the overall protein length. The black arrow highlights the characteristic DciA domain.

#### Supplemental Tables

[The data tables are available in tsv and xslx formats, and on GitHub]

**Table S1. Bacterial DciA domain-containing proteins.** Protein, domain architecture, group classification, DciA and IDR domain length, and lineage-related metadata for all DciA domain-containing proteins. The table legend carries detailed explanation of each column in the data file.

**Table S2. Archaeal and Eukaryotic DciA domain-containing proteins.** Protein, domain architecture, group, and lineage-related metadata for each of the archaeal and eukaryotic proteins are shown in this table. See table legend for Table S1.
